# Global review of meta-analyses reveals key data gaps in agricultural impact studies on biodiversity in croplands

**DOI:** 10.1101/2024.04.19.590051

**Authors:** Jonathan Bonfanti, Joseph Langridge, A. Avadí, N. Casajus, A. Chaudhary, G. Damour, N. Estrada-Carmona, S. K. Jones, D. Makowski, M. Mitchell, R. Seppelt, Damien Beillouin

## Abstract

**Aim:** Agriculture depends heavily on biodiversity, yet unsustainable management practices continue to affect a wide range of organisms and ecosystems at unprecedented levels worldwide. Addressing the global challenge of biodiversity loss requires access to consolidated knowledge across management practices, spatial levels, and taxonomic groups.

**Location:** Global

**Time period:** 1994 to 2022

**Major taxa studied:** Animals, microorganisms, plants.

**Methods:** We conducted a comprehensive literature review synthesising data from all meta-analyses about the impacts of agricultural management practices on biodiversity in croplands, covering field, farm, and landscape levels. From 200 retained meta-analyses, we extracted 1,885 mean effect sizes (from 69,850 comparisons between a control and treatment) assessing the impact of management practices on biodiversity, alongside characterising over 9,000 primary papers.

**Results:** Seven high-income countries, notably the USA, China, and Brazil dominate agricultural impact studies with fertiliser use, phytosanitary interventions and crop diversification receiving widespread attention. The focus on individual practices overshadows research at the farm and landscape level. Taxonomically, Animalia, especially arthropods, are heavily studied while taxa such as annelids and plants receive comparatively less attention. Effect sizes are predominantly calculated from averaged abundance data. Significant gaps persist in terms of studies on the effects of agricultural interventions on specific taxonomic groups (e.g. annelids, mammals) and studies analysing functional traits.

**Main conclusions:** Our study highlights the importance of analysing the effects of combined practices to accurately reflect real-world farming contexts. While abundance metrics are common, reflecting several biodiversity facets and adopting a more balanced research approach across taxa are crucial for understanding biodiversity responses to agricultural changes and informing conservation strategies. Given the unbalanced evidence on impacts of agricultural practices on biodiversity, caution is required when utilising meta-analytical findings for informing public policies or integrating them into global assessment models like life-cycle assessments or global flux models.

## 1. Introduction

Human alteration of the Earth’s land surface is substantial and growing (Winkler *et al*. 2021). Therefore, managing sustainably agricultural land - the world’s largest managed biome - is crucial for preserving biodiversity and its vital contributions to people at local and global scales (Campbell *et al*. 2017; IPBES 2019). While conservation-focused practices such as habitat preservation and low-input approaches generally yield positive biodiversity impacts (Beillouin *et al*. 2021; Estrada-Carmona *et al*. 2022; Newbold *et al*. 2015; Tamburini *et al*. 2020), evidence suggests that unsustainable agricultural management practices continue to threaten the long-term conservation of various species globally, in particular, vertebrates (Rigal *et al*. 2023), invertebrates (Raven & Wagner 2021), and plant species (Biesmeijer *et al*. 2006; Dorrough & Scroggie 2008). Agriculture’s negative impacts can occur at localised or broader scales and may vary depending on the species, biome, and specific practices employed (Cozim-Melges *et al*. 2024; Rocha-Ortega *et al*. 2021; Troudet *et al*. 2017). Ensuring that areas under agriculture are managed through the sustainable use of biodiversity and innovative agroecological approaches (in line with Target 10 of the Global Biodiversity Framework - GBF) is a major societal challenge. Actions planned to achieve the GBF targets should be based on robust and comprehensive scientific evidence.

The adoption (or non-adoption) of agricultural practices is greatly influenced by market and non-market incentives across public and private policies, but these incentives are often not effective at enhancing environmental outcomes (Piñeiro *et al*. 2020). Quantifying the dynamics of land-use change and management practices across scales is critical in order to enable the design of incentives effective at halting biodiversity loss and restoring ecosystem functioning. This requires precise quantitative data on the impacts of a wide range of agricultural management practices. But with thousands of primary research papers published globally each year, expecting policy-makers and farmers to digest such extensive information to enhance biodiversity outcomes and agricultural performance may be unrealistic. Parallelly, this vast body of primary knowledge in agricultural topics has led to a surge in first-order meta-analyses in recent years (e.g. Beckmann *et al*. 2019; Estrada-Carmona *et al*. 2022; Jones *et al*. 2023). Policy decisions would benefit from prioritising meta-analyses, as they offer expedited access to synthesised findings from a significant body of literature on specific topics (Kavale & Forness 2000; Miteva *et al*. 2012). Yet this exponential growth of syntheses inherently increases the risk of selection bias, or picking certain types of evidence syntheses over others (Haddaway *et al*. 2020). Equally, the diverse, even sometimes contradictory, results across meta-analyses could make it difficult to design and implement effective environmental policies. Using systematic approaches to consolidate a large body of the existing research into multi-level reviews may be important for providing a clear and precise examination of the specific types of agricultural management, studied species taxa, and potential moderator effects (Dainese *et al*. 2019; Davison *et al*. 2021; Garibaldi *et al*. 2019).

Although previous global syntheses exist on specific agricultural topics (e.g. Babin *et al*. 2023; Beillouin *et al*. 2021; Cozim-Melges *et al*. 2024), to our knowledge there is a distinct lack of state-of-the-art literature reviews regarding cropland management impacts on biodiversity across scales. Accordingly, we systematically review and characterise the evidence provided in 200 meta-analyses (synthesising data from over 9000 unique primary research papers) concerning the effects of agricultural and land management practices on biodiversity in croplands worldwide. Our main objective was to characterise the available evidence on agricultural impacts on biodiversity across scales and identify persistent research gaps. We use this to provide recommendations for future research and science-policy integration, to accelerate evidence-based solutions and decision-making in the transition to biodiversity-friendly croplands worldwide.

## 2. Material and Methods

### 2.1 Literature search and study selection

We conducted a comprehensive systematic literature search in Web of Science Core Collection (WOS), Scopus, Ovid, and Google Scholar to retrieve peer-reviewed meta-analyses on the impacts of agriculture on biodiversity published in English or French, in September 2022 (see PICOC framework used in Table S1, the search strings used in Table S2). The search yielded 4,154 records after removing duplicates (Figure S1).

All agricultural management practices (i.e. interventions) impacting biodiversity (i.e. study populations) at the field, farm or landscape level (i.e. context) were retained (for intervention definitions, see Table 1). All relevant biodiversity metrics (e.g. abundance, biomass, richness), from which the impacts of an intervention can be reliably demonstrated were retained. Lastly, all effect size indices (e.g. odds ratio, Hedges’ g, Cohen’s d, *etc.*) that quantify the magnitude and direction of the effect of an intervention on the study population were retained.

**Table 1.** A list of agricultural management definitions used in the review process for post-classifying interventions.

| Intervention group | Intervention | Definition(s) |
| --- | --- | --- |
| Individual practice | Agroforestry | A farming practice in which at least one woody perennial (e.g., trees, shrubs) species is deliberately grown on the same land-management units as crops (Beillouin et al. 2019). |
|  | Crop diversification | Agricultural practices leading to increasing crop diversity in a field in space and/or time. This could include longer rotations, addition of cover crops, multiple cropping, cultivar mixtures, or intercropping (Beillouin et al. 2019). |
|  | Fertilisers and Soil amendments | Methods used to improve the supply of essential elements (mainly nitrogen, phosphorus, and potassium) to plant growth by chemical or organic external inputs (FAO 2024). |
|  | GMO | Genetically modified organisms, here a crop in which one or more changes have been made to the genome, typically using high-tech genetic engineering (e.g. Bt - *Bacillus thuringiensis* - maize) (FAO 2024). |
|  | Pest and disease management | Any chemical, physical, or other type of agricultural practice that aims to directly prevent pests and diseases from affecting a crop (FAO 2024). |
|  | Residue management | Any agricultural practice that relates to the fate of materials remaining in the field after crop harvest (e.g. residues retention) (FAO 2024). |
|  | Tillage management | Any soil mechanical cultivation practice (e.g. low-till) (FAO 2024). |
|  | Water management | Practices used for maintaining crop water supply. |
| Agricultural system | Combined practices | Combination of at least two individual practices used simultaneously in the same field. |
|  | Conservation agriculture | A farming system based on the simultaneous maintenance of a permanent soil cover, minimum soil disturbance, and diversification of plant species (FAO 2024). |
|  | Organic agriculture | A farming system defined by official standards based on agricultural practices which, among others, exclude the use of synthetic biocides and fertilisers, and the use of genetically modified organisms (GMOs) (FIBL & Forschungsinstitut für biologischen Landbau 2021). |
| Landscape | Landscape complexity management | Specific landscape patterns and characteristics encompassing configuration, heterogeneity, or composition of an agricultural landscape (Estrada-Carmona et al. 2022). |
|  | Land-use change | Any modification of the purpose or function for which a particular area of land is used over time. Here, any conversion from cropland to any other land (e.g. forest land, grassland, wetlands, settlements) and vice-versa (IPCC 2003). |

We undertook a two-stage screening process on all records in Abstrackr (Wallace *et al*. 2012). Firstly, article titles and abstracts were screened and any clearly illegible article removed. Secondly, the full texts of all remaining articles were screened. Articles were retained based on the following eligibility criteria: (i) the article is a meta-analysis, i.e. it presents a quantitative synthesis of experimental results (paired-data) retrieved from several primary studies, (ii) the meta-analysis quantifies the effect of any agricultural management practice in croplands on biodiversity, (iii) the article is written in English or French. Lastly, no date or geographical restriction were applied. The final database, its structure as well as the systematic review methodology were extensively described in Bonfanti *et al*. (2023).

### 2.2 Data coding and evidence mapping

We characterised the evidence based on the number of i) meta-analyses, ii) effect sizes (i.e. overall estimate of an intervention’s impact on a specific outcome compared to a control, derived from multiple primary studies), iii) paired-data (i.e. comparative outcome data obtained from a matched control and treatment group), and iv) primary studies (Figure 1).

**Figure 1.**
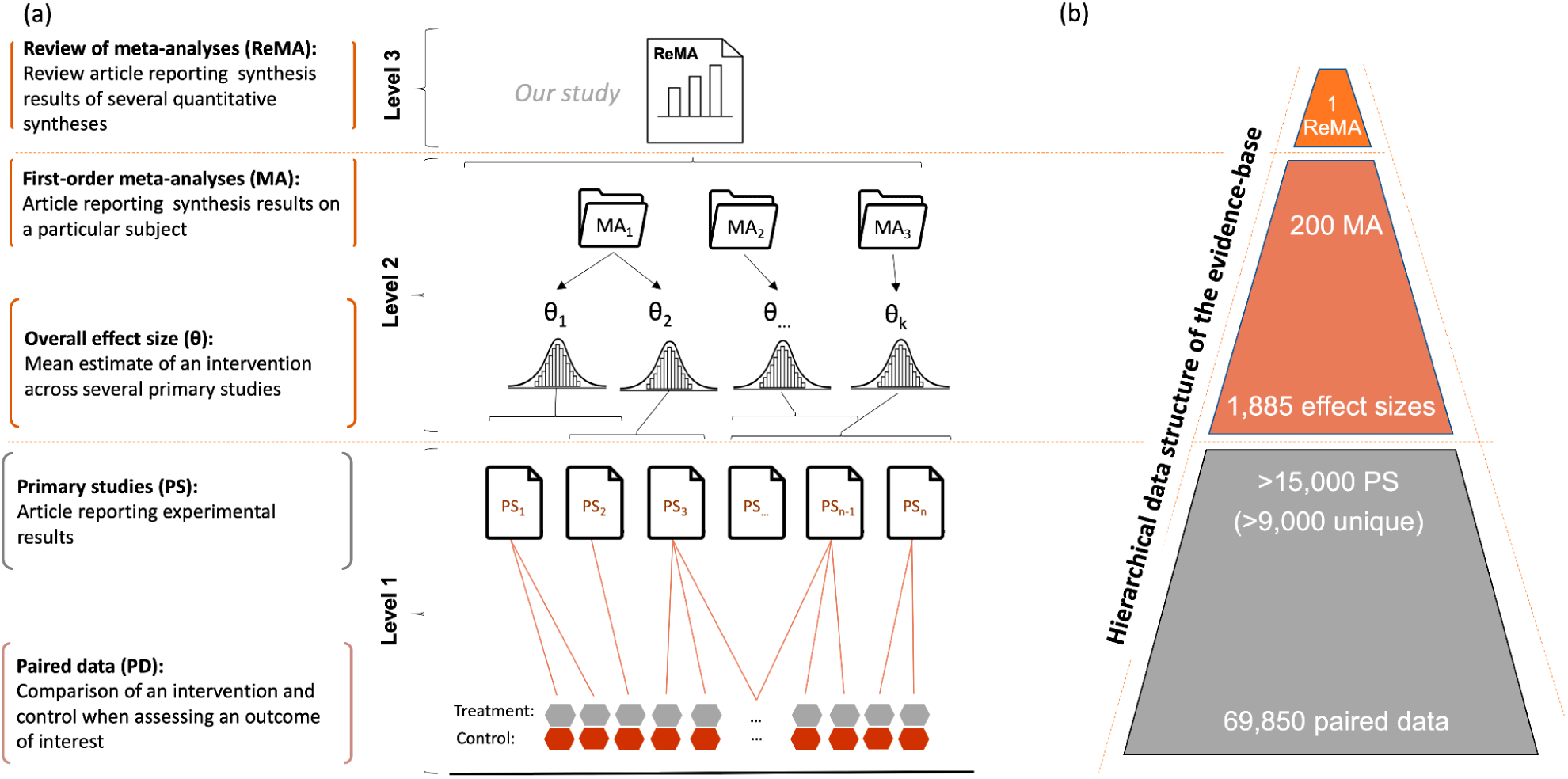
A conceptual representation of the multi-level nature of the evidence (a) and illustration of the present review (b). At the lowest level (level 1), the individual primary studies report one or several paired data comparisons on an outcome of interest. At level 2, meta-analyses report one or several mean estimates (i.e. effect-size) of an intervention across several primary studies. Level 3 is a descriptive state of the art (i.e. an evidence map) of 200 meta-analyses quantifying a measure of impact of an agricultural practice on biodiversity. These meta-analyses presented 1,885 effect-sizes from 69,850 paired data reported in >15,000 total primary studies, of which >9,000 primary studies are unique (i.e. one primary study could be used in several meta-analyses).

We extracted and assigned the highest available taxonomic resolution to each effect size based on the information in the retained meta-analyses, and using the Catalogue of Life classification (COL, Bánki *et al*. 2024). We thus mapped study populations first to 1) kingdom (e.g. *Animalia*), 2) phylum (e.g. *Arthropoda*), 3) class (e.g. *Insecta*), and 4) order level (e.g. *Hymenoptera*). When authors mentioned an ecological group *sensu lato* (e.g. feeding guilds within nematodes), this information was coded. We compared taxonomic group occurrences in our dataset against those reported in biodiversity-focused scientific literature reported in a randomly-sampled subset (*ca*. 10%) of all articles listed in Web of Science using the word ‘biodiversity’ in their title, provided by Mammola *et al*. (2023). To do so, we mapped Mammola *et al*. (2023) occurrences to a comparable taxonomic resolution (according to COL) i.e. at a kingdom level for microorganisms and plants, and at a phylum level for animals.

Each effect size was assigned a biodiversity metric: abundance, activity, biomass, phylogenetic diversity, taxonomic diversity, taxonomic richness, trait-based, or ‘multiple metric’. If a meta-analysis quantified the biodiversity response using more than one metric, we extracted data for each metric. When a meta-analysis presented subgroup analyses by taxa or other moderators (e.g. climate type, soil parameters, etc), we extracted the effect size with the highest taxonomic resolution to avoid redundancy in our database. The interventions associated with each extracted effect size were coded and categorised into three main groups: i) individual practices, ii) agricultural systems, and iii) landscape-level practices (Table 1). We categorised the various effect size indices into six main types (Borenstein *et al*. 2009): mean difference, standardised mean difference (e.g., Hedge’s g), response ratio (e.g, Ln(RR)), odds ratio, and correlation regression (see Figure S6).

We extracted the references of all primary studies used in each meta-analysis. The unique digital object identifier (DOI) associated with each primary study was used to identify the list of unique primary studies. For each primary study, we extracted information using text-mining techniques, based on a predefined keyword list, on the study location (i.e. country) and the type of agricultural management practice. The results were then checked manually to ensure reliability of the extracted information. Country names and agricultural practices were extracted either from the title, the abstract, or the material and methods section. Agricultural practices were coded and categorised using the same method as the effect sizes (Table 1). Among the 9,080 unique primary studies, 5,600 provided enough information to code the location.

All graphical representations were performed in R environment (version 4.3.1, R Core Team 2021) notably using the packages `tidyverse` (Wickham *et al*. 2019), `ggplot2` (Wickham 2016), and in RAWGraphs (Mauri *et al*. 2017).

## 3. Results

### 3.1 High-income countries dominate primary studies of agricultural impact on biodiversity

Our review identified 200 meta-analyses published between 1994 and 2022 covering *ca*. 9,000 unique primary publications (and ca. 15,000 non-unique references) resulting in 1,885 effect sizes (Figure 1). The primary data synthesised in the meta-analyses originate from studies conducted all over the world (Figure 2), but evidence is uneven. For instance, Asia contributed 25.8% of the papers with accessible information on location, followed by Europe (24.6%), North America (20.3%), and South America (16.0%). Less than 10% of the primary studies originate from Africa and Oceania. Seven high-income countries contributed to more than 50% of the primary studies with information on locations from the USA (891 studies), China (845), Brazil (347), Canada (266), Germany (244), Australia (197), and Spain (190). Fertilisers and organic amendments emerge as the most extensively prevalent practices studied, featuring as the top two most studied practices in five out of six world regions. Crop diversification consistently ranks among the top four most studied practices, with cover crops, crop rotations, and intercropping being the main practices, comprising approximately 50%, 25%, and 7% of studies, respectively. Tillage is in the top four practices in four out of six world regions. Globally, landscape complexity receives greater attention than land-use change, and organic agriculture is more frequently studied than combined practices or conservation agriculture across regions. Our review uncovers regional specificities in agricultural research. For instance, the effects of agroforestry systems are more frequently investigated in Africa and South America than in other regions, while pest and disease management receives more attention in North America and Oceania.

**Figure 2.**
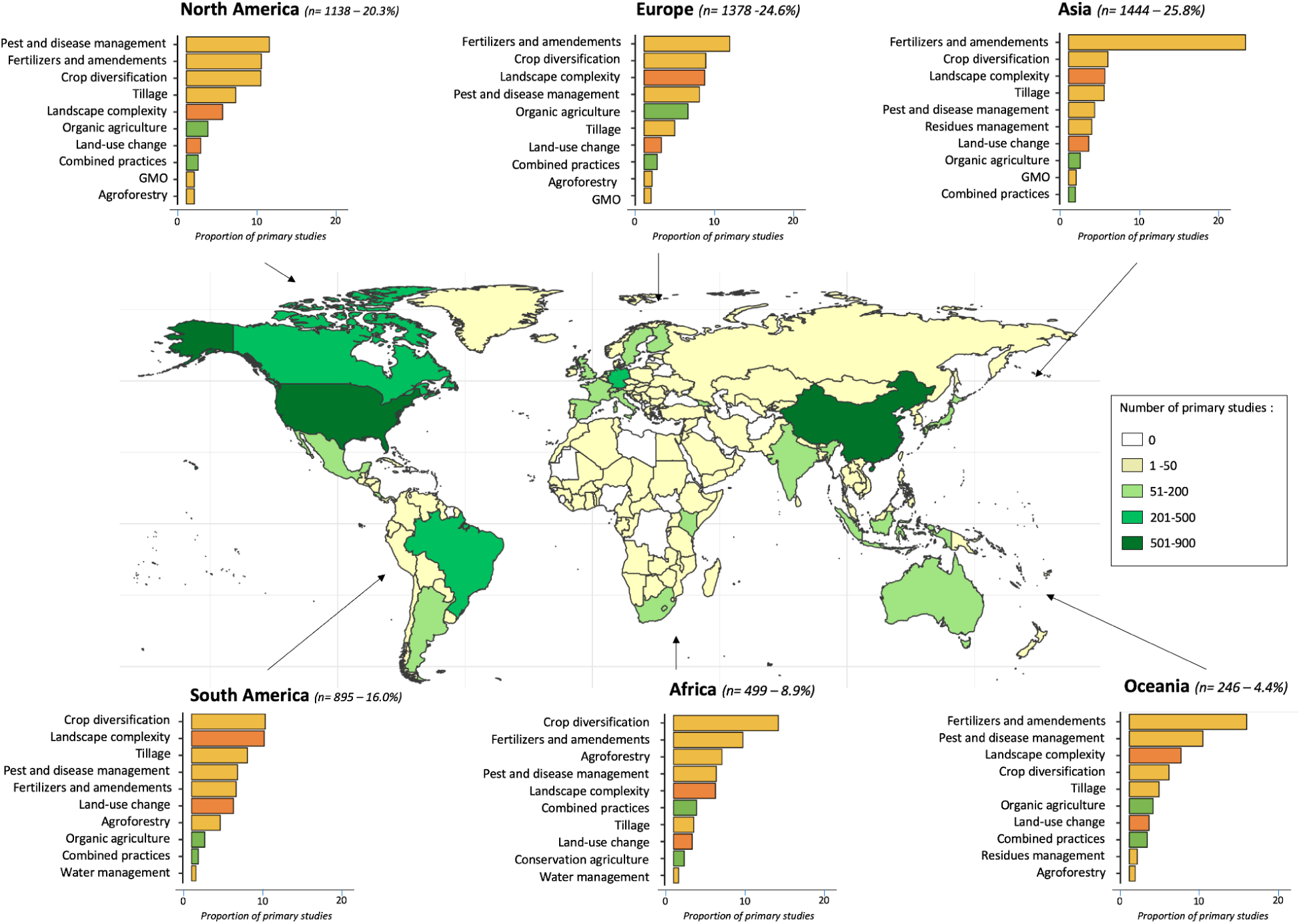
Locations of the primary studies included in the 200 meta-analyses and the top 10 agricultural interventions analysed by world region. Some meta-analyses did not provide the references of their original studies, and the full text or locations of some primary studies were not available, thus not accounted for. Among the 9,080 unique primary studies, 5,600 provided enough information to code the location. In the barplots, yellow bars: field-level practices; green bars: farm-level practices; orange bars: landscape-level practices.

### 3.2 Agricultural inputs, phytosanitary interventions, and crop diversification dominate in evidence syntheses

Our study shows that evidence from synthesis research is skewed towards studying the effect of individual practices, with these representing six times (x 6.4) more effect sizes than agricultural system-level interventions, and four times (x 4.1) more than landscape-level (Figure 3). The gap in evidence is even larger when we look at the number of primary studies rather than the effect-sizes: we observe a 11.4-fold more studies focused on individual practices compared to agricultural systems, and a 4.92-fold more compared to landscape factors. In meta-analyses focusing on individual practices, the spotlight is on practices related to inputs such as fertilisers and amendments, and residue retention (440 effect sizes in total, accounting for 32% of the effect sizes data related to individual practices). Following closely are phytosanitary interventions, including pest and disease treatments and GMOs represented by 369 effect sizes in total (27%). Practices related to on-farm diversification strategies, for example agroforestry and crop diversification, represent almost one third of the total effect sizes assessed (373 effect sizes in total; 28%). Less studied practices include tillage, represented by one-tenth of effect sizes (161 effect sizes, 12%) (Figure 2, Figure 3). Regarding landscape-level interventions, syntheses focus primarily on the effect of land-use change (273 effect sizes) and less commonly on landscape complexity (57 effect sizes). This hierarchy seems to be reversed at the level of primary studies, *i.e.* more studies focus on landscape complexity than land-use change (Figure 2). Finally, agricultural-system level practices are mainly represented by organic agriculture interventions (142 effect sizes), followed by combined practices (57 effect sizes) and conservation agriculture (11 effect sizes). Water management is scarcely addressed.

**Figure 3.**
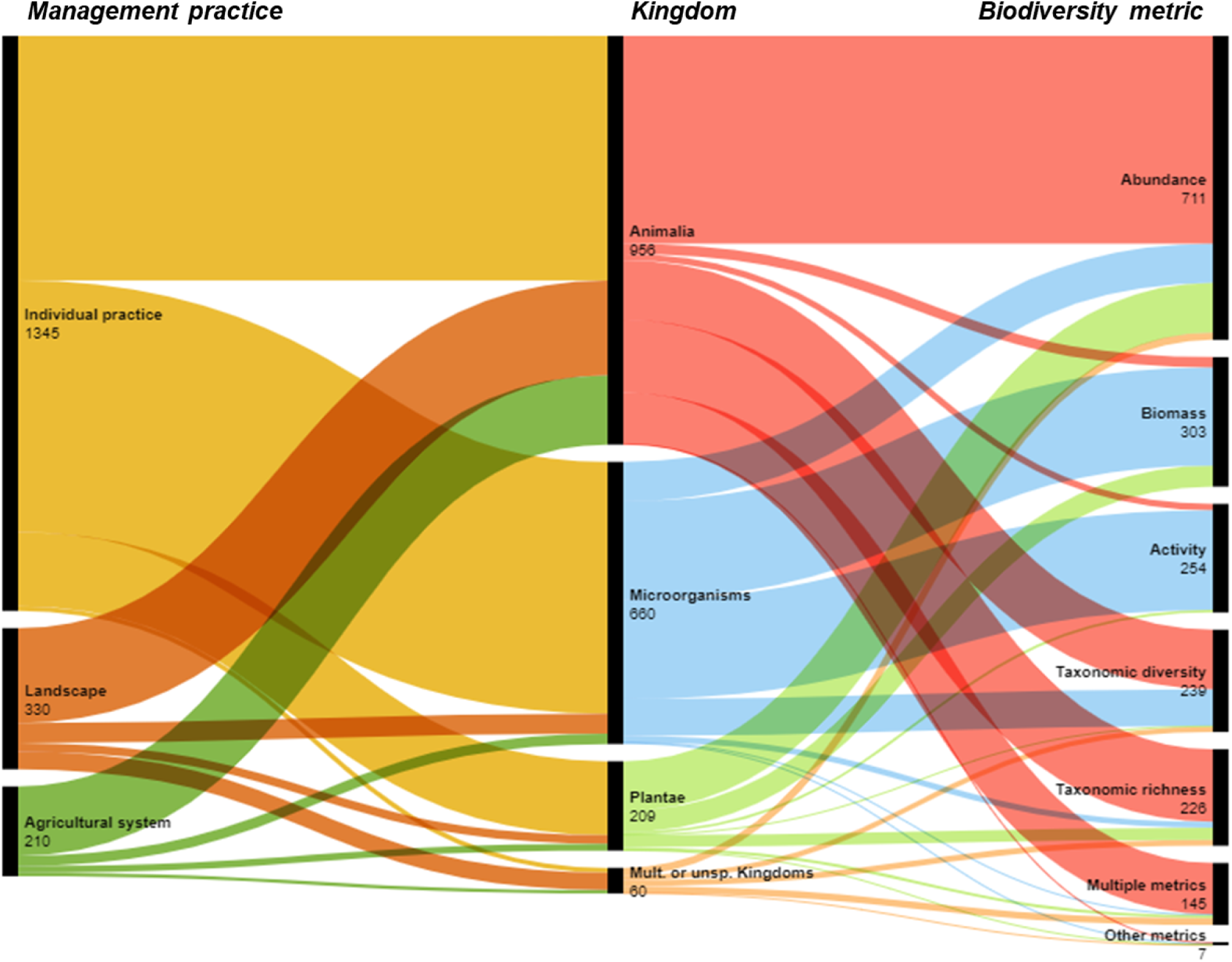
Distribution of 1,885 mean effect sizes reported in the 200 meta-analyses studying the effects of agricultural interventions on biodiversity. The types of agricultural interventions are shown on the left bar, biodiversity taxa at kingdom level are shown on the middle bar, with biodiversity metrics shown on the right bar; numbers represent effect sizes. ‘Mult. or unsp. Kingdoms’ indicates effect sizes combining information on several taxa. For clarity, on the middle bar: ‘Archaea’ (n=3) were grouped within ‘Bacteria’, and ‘Chromista’ (n=3) were grouped within ‘Fungi’. On the right bar: ‘Other metrics’ refers to ‘Phylogenetic diversity’ (n=3) or ‘Trait-based’ (n=4) metrics. ‘Multiple metrics’ indicates effect sizes aggregating multiple or unspecified biodiversity metrics.

### 3.3 Insects, birds, and abundance outcomes prevail in evidence syntheses

Synthesis research reveals a strong taxonomic bias. Animalia accounts for half of the available effect sizes (50.5%) (Table 2, Figure 3). Within this group, arthropods represent 46.5% of effect sizes, of which 61.0% are insects, 10.0% arachnids and 26.8% other or unspecified arthropods (Table 2). Nematodes, Vertebrata, and Annelids are significantly underrepresented compared to Arthropods, by a factor of 2.2, 4.7, and 18.5, respectively; similar trends exist for paired data. Moreover, 67% of effect sizes within vertebrates concern only birds. Among other kingdoms, Plants contribute 11.1% of all effect sizes, followed by Fungi at 5.6% and Bacteria at 4.7%, with the remaining one-quarter of the available effect-sizes (24.3%; 459 effect sizes) not assigned to a single kingdom because the data concerns several or unspecified taxa (Table 2).

**Table 2.**
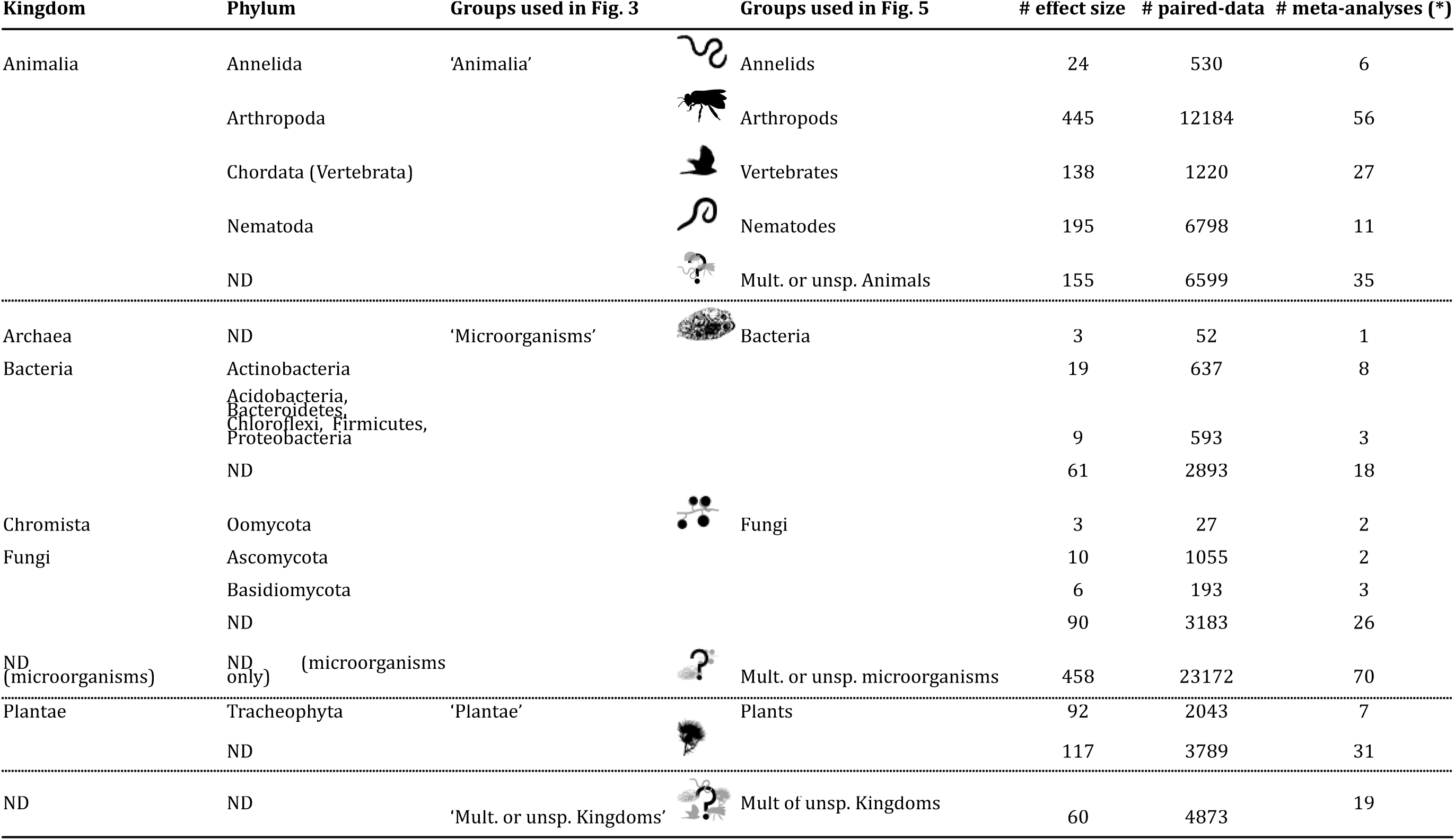
Distribution of evidence in terms of taxonomic structure (Kingdom and Phylum levels). Mult.or.unsp.: multiple or unspecified, i.e. some effect sizes could not be attributed to a single taxonomic group. (*) One meta-analysis may provide several effect sizes belonging to several taxonomic groups

The taxonomic resolution of the effect sizes largely differs between Kingdoms (Figure 4). Microorganisms had low taxonomic resolution, often summarised through non taxa-specific metrics (e.g., microbial biomass carbon). When Bacteria or Fungi kingdoms were identified, few effect sizes were characterised at a higher resolution. Taxonomic resolution of plants was also generally low, with 44% effect sizes classified at phylum level, and the majority of all plants falling under the ‘weeds’ functional group designation (80%) (Figure S4). Animalia was the most detailed kingdom, with 84% of effect sizes detailed at phylum, 51% at class, and 32% at order level. Note that for animals, when the taxonomic resolution was very low (i.e. kingdom level), an ecological group was given in 76% of the effect sizes (e.g. ‘feeding guilds’ in different groups, ‘functional groups’ such as pollinators, natural enemies, etc.).

**Figure 4.**
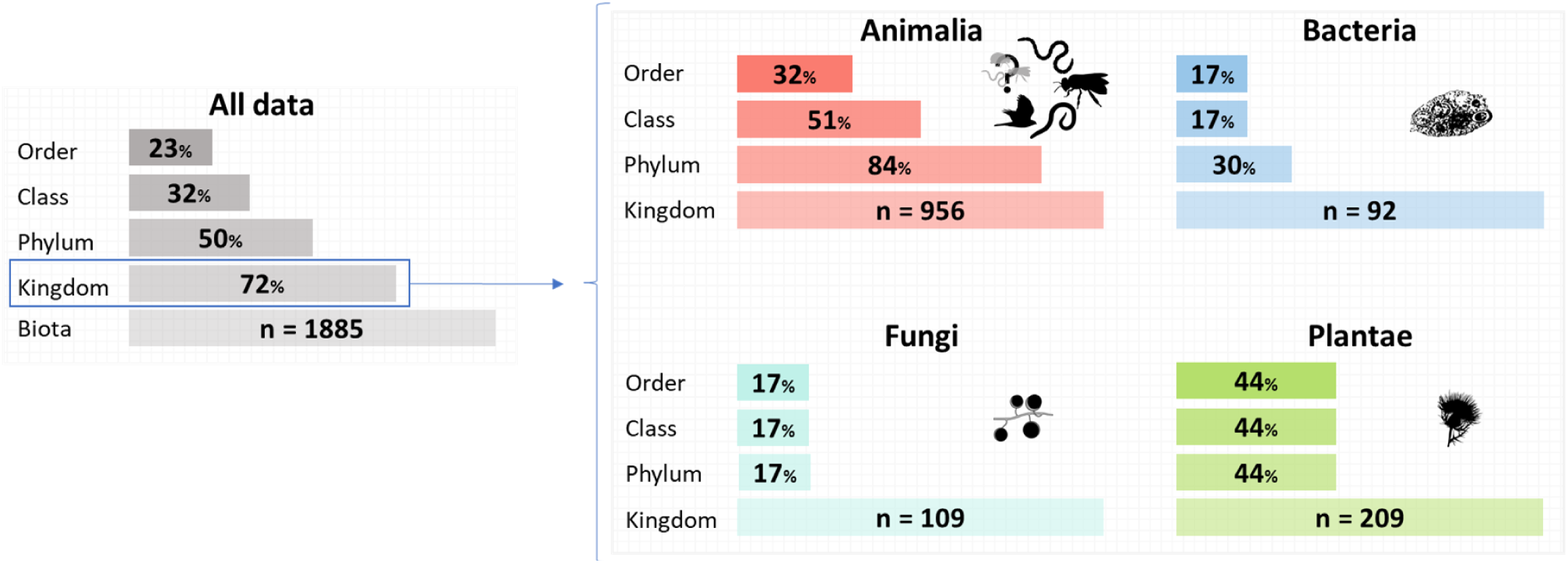
Taxonomic resolution for each major Kingdom and for the total 1,885 effect size data. The total number of effect sizes per Kingdom is depicted at the bottom of each plot, while the available information at lower taxonomic ranks is represented as a percentage of the total effect sizes.

Abundance was the most frequently used metric for biodiversity, especially for plants, unspecified or multiple invertebrates, nematodes, and insects (Figures 3, Figure 5). Biomass (16%) and activity (13%) metrics were dominant for microorganisms. Taxonomic diversity and richness were less frequently used (13% and 12%, respectively), but were common for animal taxa, like insects and nematodes. Phylogenetic diversity and trait-based indices were rarely employed (<1% each) in the retained meta-analyses.

**Figure 5.**
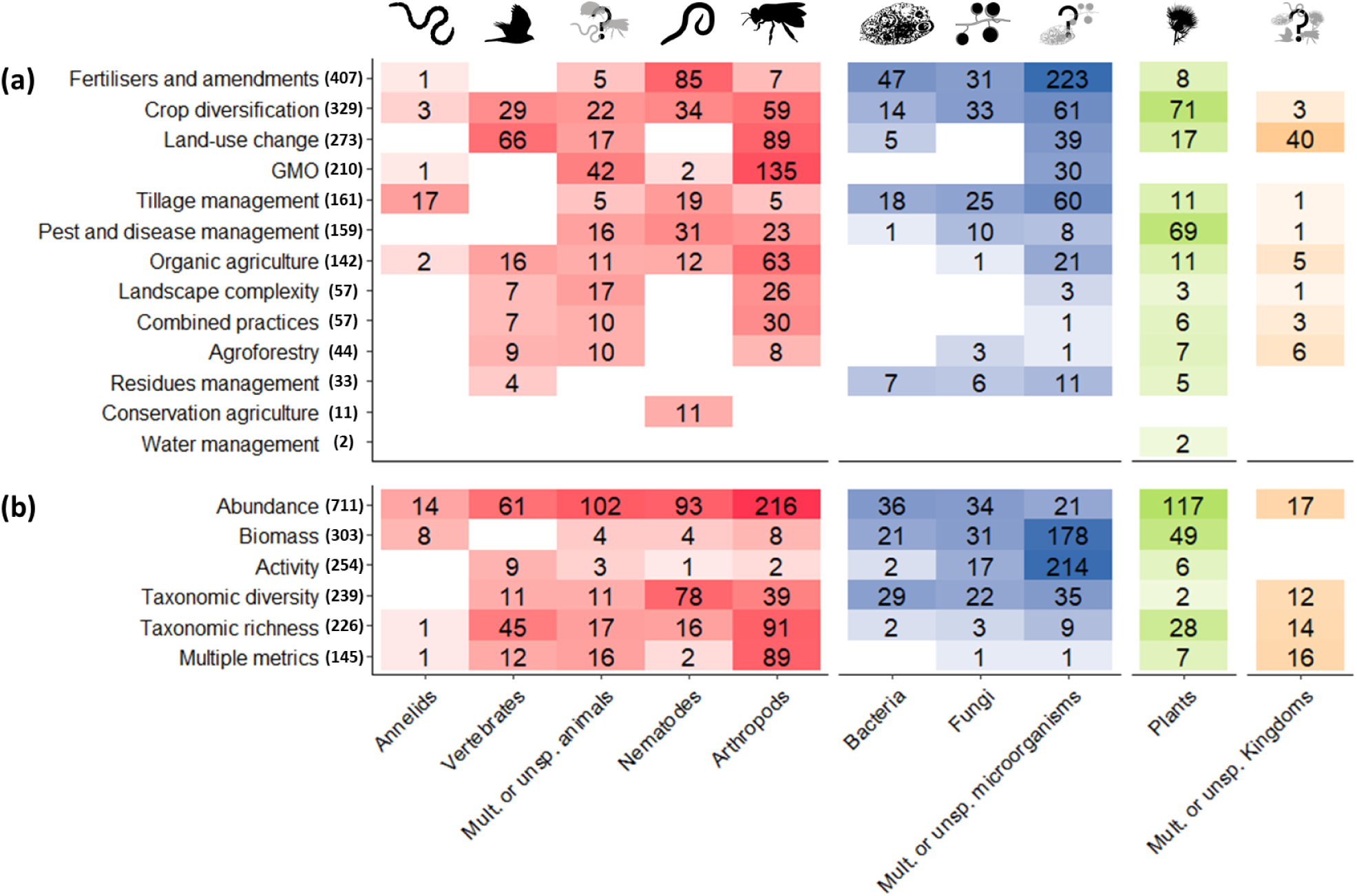
Number of effect sizes extracted from the 200 meta-analyses by taxonomic group and agricultural management practice (a), and by biodiversity metric (b). Tile labels and colour intensity represent the number of effect sizes. Taxonomic groups are presented at Kingdom level, except animals at Phylum level (see Table 2). For clarity, Archaea are grouped with Bacteria, Chromista are grouped with Fungi, and effect sizes depicting phylogenetic and trait-based metrics (n=7 in total) are not shown. The number in parentheses on the x-axis represents the total number of effect sizes for each management practice and biodiversity metric. Tile colours: red for animals, blue for microorganisms, green for plants, orange for multiple or unspecified kingdoms.

In addition, our results on taxa occurrences align somewhat with global biodiversity patterns i.e. Mammola *et al*. (2023) concerning animals in general (Figure 6). However, within animals, arthropods and nematodes are over-represented in syntheses on biodiversity in agroecosystems (our database) by approximately a factor of 2 and 10 respectively, while vertebrates are under-represented by a factor 2 relative to Mammola *et al*. (2023). In general, microorganisms are over-represented in agrobiodiversity syntheses (x3) and plants receive much less focus (4 times less). Other taxa such as Mollusca, Platyhelminthes, and Protozoa are absent from our dataset but are the focus of respectively 0.7, 0.1 and 0.1 % of the studies in the general biodiversity literature; Chromista is characterised in 3 effect-sizes in our dataset but absent in Mammola *et al*. (2023).

**Figure 6.**
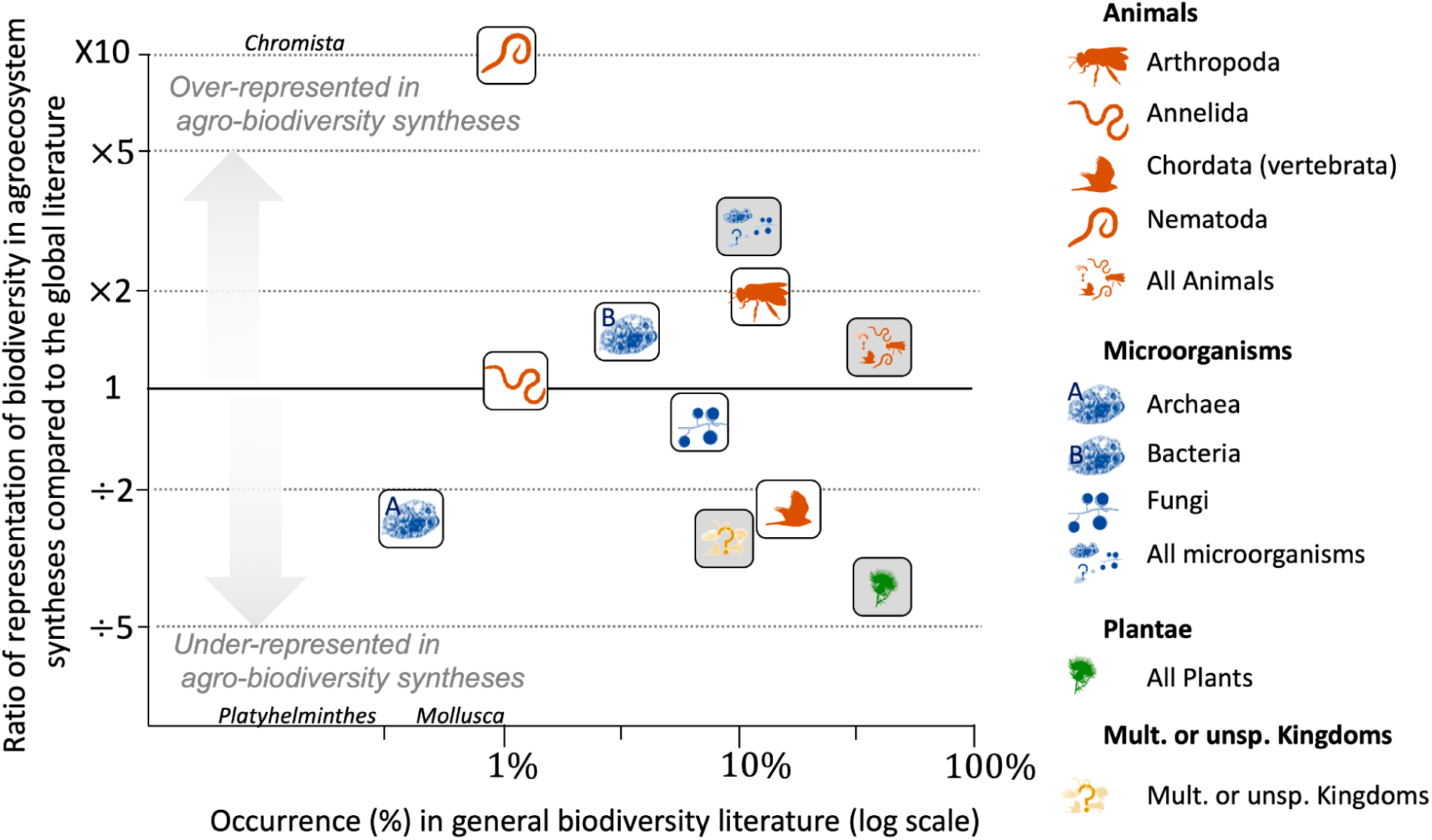
Over- or under-representation of taxonomic groups in syntheses on biodiversity in agroecosystems (our study) compared to the general biodiversity literature (data from Mammola *et al*. 2023). The y-axis represents the ratio between the percentage of occurrence of each taxonomic group in both studies, the x-axis represents the percentage of occurrence of taxonomic groups in Mammola *et al*. (2023) after harmonisation with respect to the Catalogue of Life classification. The four major groups (see Figure 3, Table 2) are presented with grey background boxes, and detail is given with white background boxes. Microorganisms are detailed at Kingdom level and animals at Phylum level. When no box is drawn, the group is absent from one of both studies, i.e. Chromista is absent from Mammola *et al*. (2023), Mollusca and Platyhelminthes are absent from our study. For clarity, Protozoa and Algae are hidden behind Platyhelminthes and Mollusca, respectively. Raw data are given in Table S3.

### 3.4. At least half of the evidence is missing on agricultural intervention impacts on biodiversity

Combining results on agricultural practices and biodiversity outcomes, our study reveals that 47 out of 130 theoretically possible combinations between agricultural interventions effects on specific taxonomic groups have not been studied in any meta-analysis, while 18 combinations are filled with three or fewer effect sizes (Figure 5). Most effect-sizes represent the same intervention-outcome combinations, e.g. the effect of fertilisers & amendments on microorganisms, crop diversification on plants and microorganisms, or genetically modified organisms on insects (Figure 5). As a result, almost no information is available on some potentially important agricultural drivers of biodiversity loss, e.g. effect of land-use change on underground biodiversity, or of crop diversification, tillage, agroforestry, organic agriculture on herptiles, mammals. Water management impacts on all taxa are considerably understudied.

## 4. Discussion

The 200 meta-analyses identified in this study on the effect of agricultural practices on biodiversity could serve as a resource in addressing the urgent need to slow agricultural-driven biodiversity loss. Aligning agricultural research efforts with growing demand for evidence-based policymaking is a gateway to finding and implementing more biodiversity-friendly agricultural systems (Semenchuk *et al*. 2022; Sutherland *et al*. 2004). This scientific literature can provide insights on how to re-design heavily intensified agricultural systems and landscapes toward biodiversity-friendly agriculture (e.g. Stein-Bachinger *et al*. 2022) notably in the world’s top agricultural producers and exporters where most of the data originates (i.e. USA, Brazil, China, Canada and France). It may also allow for constructing objective arguments to conserve traditional or highly biodiversity-friendly practices (Herrero *et al*. 2017; Hutchins *et al*. 2024) but which face political and economic pressure in many world regions especially tropical and subtropical - to transition towards conventional and intensified systems (e.g. Perfecto *et al*. 1996; Tscharntke *et al*. 2012). Yet, our systematic review reveals gaps in knowledge that may hinder evidence-based decision making for certain regions, intervention scales, and taxa. Unbalanced knowledge and persistent research gaps on the relation between agricultural practices, ecological processes, and ecosystem services, coupled with the absence of adapted and/or localised knowledge, constitute two primary factors that constrain the practical application of biodiversity-based agriculture (Duru *et al*. 2015).

We therefore identify five shortcomings to be addressed through future research:

1. The reductionist approach dominant in studies of the effects of agricultural practices on biodiversity needs complementing with systems approaches better suited to understanding the complexities of farming systems. Specifically, research is needed across multiple spatial levels beyond individual practices in isolated fields. Understanding the performance of individual practices is important to manage local impacts on biodiversity, yet comprehending whole farm and landscape level impacts is paramount for effectively managing agricultural systems so people can live in harmony with nature in line with the Global Biodiversity Framework’s post-2020 goals. Given that farmers seldom adopt singular practices in isolation, more studies monitoring the outcomes of combined practices (e.g. intercropping with cover crops and organic amendments) are needed to adequately reflect the complexity and dynamics of real-world agricultural systems. In our database, system-level agricultural land management (e.g. conservation agriculture) remains heavily underassessed in comparison to some individual practices (e.g. fertilisation alone). Combining multiple practices or implementing integrated agricultural systems generates more positive outcomes (e.g. Beillouin *et al*. 2019; Rosa-Schleich *et al*. 2019). More research to understand which practice bundles are most effective would help minimise potential trade-offs to biodiversity or food production (Jones et al. 2023). Additionally, the paucity of quantification on the effects of landscape-level interventions in certain regions (e.g. land use change in Africa) needs addressing to enable more effective design of integrated landscape approaches, ideally as part of multi-level research designs that capture the effects of field to landscape management decisions. Indeed, both field-level and landscape-level interventions are essential to restoring biodiversity and the two scales are interdependent (Estrada-Carmona *et al*. 2022; Lichtenberg *et al*. 2017). The present review challenges the notion that agricultural effects are thoroughly studied (e.g. Ortiz *et al*. 2021). Advancing the relationship between agricultural practices and biodiversity needs more than just additional studies, but also more effectively accounting for actual farmer practices, ultimately aligning policies with farmers’ needs or current practices (Tyllianakis & Martin-Ortega 2021).
2. Emphasis on practices aiming to increase the efficiency or substitution of inputs could limit innovation and whole-system performance assessments and support the *status quo* of agricultural practices (i.e. conventional agriculture). The dominance of studies on the impact of fertilisers in croplands is unsurprising due to their potential for securing global food security and improving food security through increased crop yields and productivity for certain crops (e.g. cereals - Falconnier *et al*. 2023). Yet, heavy use of fertilisers has considerably altered the geo-physical global cycles of major nutrients (Penuelas *et al*. 2023) putting pressure on planetary boundaries (Rockström *et al*. 2017). The ecological restoration of agricultural landscapes, along with the emulation of natural principles in agricultural practices is increasingly advocated to enhance system performance (Wezel *et al*. 2020). This shift from prioritising efficiency to improving habitat quality may necessitate different or more intricate experimental designs and research focusing on cascading effects and on the various facets of biodiversity. For instance, reducing soil tillage to enhance soil health can lead to improved soil structure and fertility, thereby reducing inputs (Willekens *et al*. 2014). This transition is already underway in scientific research, as evidenced in our study by the growing body of knowledge on diverse forms of crop diversification. Yet our analysis of primary studies suggests that the majority of crop diversification strategies primarily focus on cover crops. While these require less extensive system redesign and yield positive outcomes for ecosystem services, other diversification strategies may have more significant positive effects (Beillouin et al., 2021).
3. Agriculture impacts on a wide range of species, yet research remains focused on few taxa. Our analysis highlights taxonomic biases in the existing agricultural evidence (Troudet *et al*. 2017, Davison *et al*. 2021). For instance, while nematodes and earthworms are extensively studied in relation to tillage practices in our database, other soil fauna taxa and vertebrates receive comparatively less attention, despite their crucial role in microhabitat modifications (Rosenberg *et al*. 2023). Similarly, pest and disease management effects are predominantly studied in relation to targeted pest taxa, neglecting their broader impact on untargeted taxa (Anthony *et al*. 2023). Furthermore, plants are predominantly studied at a low taxonomic resolution and under the angle of ‘weeds’, thus overlooking the broader implications for untargeted plant species. These biases have significant drawbacks, potentially excluding rare species and leading to an excessive focus on charismatic ones (Enquist *et al*. 2019). While charismatic species may garner greater attention, their effectiveness as umbrella species in conservation efforts is debated (Davison *et al*. 2021; Simberloff 1998; Williams & Araéjo 2000) underlining the need for a more balanced research approach across taxa. Primary research should also contribute to fill key knowledge gaps related to the role and contributions of underrepresented taxa like fungi, earthworms, molluscs, and vertebrates (notably mammals), which are vital for cropland ecosystems.
4. Emphasis on abundance metrics tells only a partial story. A wide range of metrics to quantify and monitor biodiversity exist (e.g. Magurran 2013). In spite of that we found an overdependence on the abundance metric in agricultural impact studies. Indeed, this metric may certainly be appealing because it effectively reflects the decline - or recovery - of a taxonomic group (Santini *et al*. 2017). Likewise, abundance is often used to study functional groups important for agriculture such as pests and pathogens, natural enemies of pests, or pollinators. Furthermore, using the abundance metric may reduce time and cost since it can be applied to broad taxonomic groups and doesn’t require huge effort in terms of taxonomic identification, thus allowing a wider use of this indicator by non-specialists.Yet other metrics may be central to understanding the responses of different biodiversity facets to changes in agricultural systems, notably metrics at the community level. For example, functional richness can better indicate ecosystem function and stability and is sensitive to community-level changes (Lamb *et al*. 2009; McGill *et al*. 2006). Likewise, phylogenetic diversity metrics are promoted for capturing ecosystem functions and interlinkages with crop performance (Grab *et al*. 2019). Therefore, measuring the performance of agricultural systems that actively contribute to maintaining “life on earth” (SDG15) needs more comprehensive use of metrics including those that capture impacts at the species and the community level.
5. Building multifunctional agricultural landscapes requires reviewing evidence in an objective and meaningful way to better guide decision-makers and research agendas. As the body of synthesis papers published globally each year rises, it is becoming increasingly unrealistic to digest such extensive and complex information. Our global multi-level evidence map is a first step to help summarise such vast empirical evidence in relation to biodiversity. A second step is to quantitatively analyse the impacts of agricultural practices on biodiversity (i.e. in a second order meta-analyses). A third step could involve expanding the database to include multiple objectives such as yields, profitability, and climate mitigation. Providing decision-makers with timely access to scientific evidence can be facilitated by continually updating existing evidence syntheses. Such living reviews offer a pertinent strategy for bridging the gap between science and action and would benefit from cross-institute collaboration to mobilise and utilise resources from across evidence synthesis research teams to streamline efforts. Such approaches could aid policy-makers looking to inform their decisions with up-to-date, reliable evidence (Martin *et al*. 2023). Further, this could facilitate communication and collaboration between scientists and agricultural stakeholders through, e.g. evidence-based platforms, which are essential for fostering open dialogues and promoting more targeted conservation schemes (Maas *et al*. 2021). These flexible syntheses could be complementary to other global tools assessing the impact of policy or management practices such as life cycle assessments (LCA) (Leclère *et al*. 2020), and footprint analyses, which often rely themselves on parameters derived from the scientific literature.

## 5. Conclusion

Our work represents a comprehensive multi-level evidence map of quantitative syntheses about agricultural impact studies on biodiversity, at the global scale. We provide a characterisation of 200 meta-analyses and of 1,885 pooled experimental results (effect sizes) and over 9,000 unique primary studies assessing farming practices at the field, farm, and landscape level. Innovating, re-designing, and valuing agricultural farms and landscapes to positively contribute to biodiversity conservation while remaining resilient to changing conditions needs more robust scientific evidence. More than 27% of the estimated effect sizes focused on chemical fertilisers, GMO, or chemical pest and diseases management whereas conservation-oriented and sustainable practices such as conservation agriculture, agroforestry, water management are far less covered in the synthesis literature, despite their potential for achieving win-win outcomes (Jones *et al*. 2023). A large part of the evidence is focused on animals (51%) and mostly the impact of an intervention on animal and plant biodiversity is measured with the abundance metric (52%), whereas richness and diversity metrics, *a fortiori* on the functional and phylogenetic facets, were much less used despite their benefits for describing community-level changes. We provide five recommendations to improve the coverage and utility of knowledge syntheses and primary studies for science to contribute to achieving the Global Biodiversity Framework goal of transitioning to biodiversity-friendly croplands.

## Author contributions

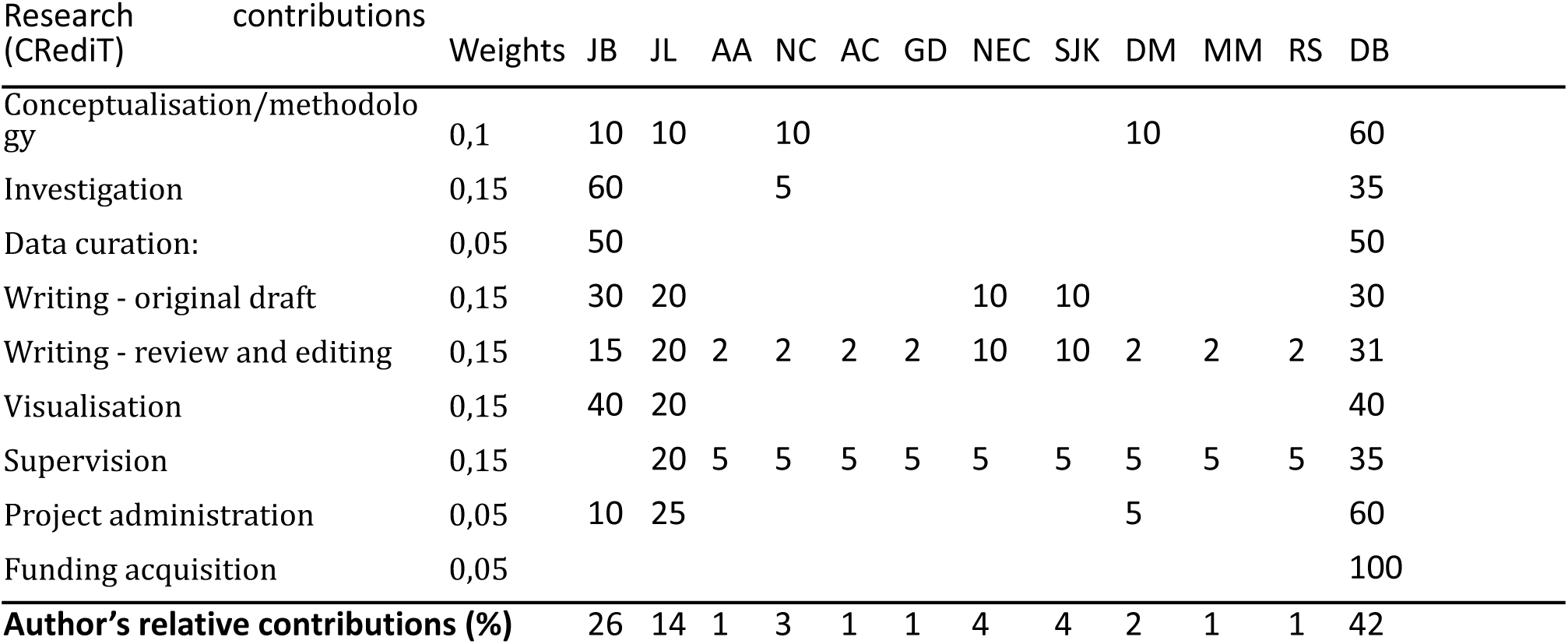

## Acknowledgements

JB is thankful to colleagues who participated in the SFE²-GfÖ-EEF International Conference on Ecological Sciences (Metz, France) for their feedback on the preliminary results of our study. JB received a postdoctoral grant from FRB (Fondation pour la Recherche sur la Biodiversité).

## Biosketch

The team’s research emphasis is primarily - but not exclusively - on the implementation and scaling of agroecology for sustainable food systems and biodiversity-friendly practices. Combined, the team has a significant number of years experience in research synthesis, aggregating evidence from multiple sources to test new conceptual hypotheses while applying quantitative and/or narrative methods to make findings from academic literature more generalisable and applicable for evidence-informed decision-making.

## Statements relating to ethics and integrity policies

## Data availability statement, accessibility

Our data consists of an Excel file database, composed of six worksheets, that is freely accessible online (https://doi.org/10.18167/DVN1/RIRTOT) and described in Bonfanti *et al*. 2023. The present paper presents results from the analysis of the ‘ES_qualitative data’ worksheet.

## Ethics approval statement

Our work did not involve live subjects (human or animal). Thus, no ethics agreement was required.

## Patient consent statement

Our research is not clinical. An informed consent document was not required.

## Permission to reproduce material from other sources

Permission was not required as material used is available under Creative Commons licences.

## Clinical trial registration

Our research is not clinical. No clinical trials registration was required.

