## Supplemental Tables and Figures for "Global review of meta-analyses reveals key data gaps in agricultural impact studies on biodiversity in croplands"

### Annexes

##### Literature search and study selection

Retrieval of all relevant peer-reviewed meta-analyses quantifying the impacts of agricultural management practices on biodiversity in croplands was based on the following PICO-C question structure (cf. Table S1):

**Table S1 - Components of the review questions.**

| PICO-C component | | Definition |
|---|---|---|
| Population: | All terrestrial and semi-aquatic taxa | This included microorganisms, vascular and nonvascular plants, invertebrates, and vertebrates. |
| Intervention | All agricultural management practices | Any individual practice (e.g. tillage, use of biocide, amendments…), set of practices or agricultural system (e.g. organic agriculture). studied at field or farm scale, or landscape metric in croplands. (cf. Table S1.3 for definitions) |
| Comparator | Two types of comparators:<br>(i) Different intensities of agricultural management;<br>(ii) Natural habitat | Concerning individual agricultural practices and the agricultural-system scale, we used the more intensified management as the control. We defined intensification as either i) an increase in the use of the external input (e.g. chemical, N fertilizer, mechanisation), or ii) a decrease in the number or evenness of cultivated plant over space or time. For the landscape scale: a decrease of landscape complexity or a change in land use. |
| Outcomes | All biodiversity metrics | Effect sizes, i.e. a weighted mean comparison between two modalities of agricultural management practices, usually an Intervention (treatment) and a Comparator (control). The effect size can average different biodiversity metrics (e.g. species richness, abundance, diversity indices…). It can be expressed in different effect size metrics (e.g. ratio, Hedge's g…) completed with indicators of precision (e.g. confidence interval). |
| Context | Type of publication | Only first order meta-analyses conducted anywhere in the world in croplands or agricultural contexts, with no temporal restriction. We did not consider vote-counting studies as meta-analyses. |

**The search string**

The literature search was undertaken on 21st July 2021 and updated in September 2022, through the following four search engines:

- Web of Science Core Collection (WOS) (Clarivate Analytics, USA),
- Scopus (Elsevier, The Netherlands),
- Ovid (Wolters Kluwer group, USA) and
- Google Scholar (Google, USA).

The search strings used for each database or search engine are given below (cf. Table S1.2):

**Table S2.** Search string used in WOS and Scopus engines.

| Search engine* | Search string |
|---|---|
| WOS, Scopus | (meta-analysis OR "systematic review" OR meta-regression OR "quantitative synthesis" OR "global synthesis" OR metaanalysis OR "quantitative review")<br><br>**AND** (crop* OR agricultur* OR farm* OR land-use OR landscape OR agroecosystem$ OR "nitrogen addition" OR "N add*")<br><br>**AND** (system$ OR practice$ OR management OR conventional OR "alternative agriculture" OR "alternative farming" OR organic OR agroecolog* OR "conservation agriculture" OR biodynam* OR permaculture OR IPM OR "integrated pest management" OR low-input OR "embedded natural" OR agroforestry OR biocontrol OR "urban agriculture" OR till* OR fertiliz* OR amendment$ OR manure OR biocide$ OR pesticide$ OR fongicide$ OR herbicide$ OR abandonment OR set-aside OR fallow$ OR "mixed crop-livestock$" OR "integrated crop-livestock$" OR "diversified crop-livestock$" OR "vegetation strip$" OR "insect strip$" OR "flower strip$" OR intensification OR diversification OR rotation OR *inter-crop* OR cover-crop* OR mixture OR "crop sequence$" OR polyculture OR semi-natural OR irrigation OR biofuel OR energy-crop$ OR simplification OR grassland$ OR "farm size$")<br><br>**AND** (biodiversity OR diversity OR richness OR abundance$ OR evenness OR divergence OR dispersion OR structure OR function OR index OR migration OR extinction OR coloni$ation OR communit* OR assemblage$ OR species OR population* OR "soil fauna" OR "soil diversity" OR "soil biota" OR "soil organism$" OR "soil biology" OR bacteria$ OR fung* OR mycorrhiza$ OR microb* OR arthropod* OR insect$ OR collembol* OR arachnid$ OR spider$ OR myriapod$ OR mollusc$ OR gasteropod$ OR annelid$ OR earthworm$ OR reptile$ OR amphibian$ OR avifauna OR bird$ OR mammal$ OR *fauna OR weed$ OR plant$ OR pollinator$ OR decomposer$ OR "ecosystem engineer$" OR pest$ OR disease$ OR "natural ennem*" OR "microb* regulator$") |

* Field tags were adapted for searches carried out in Web of Science (i.e. TS=) and Scopus (i.e. TITLE-ABS-KEY)

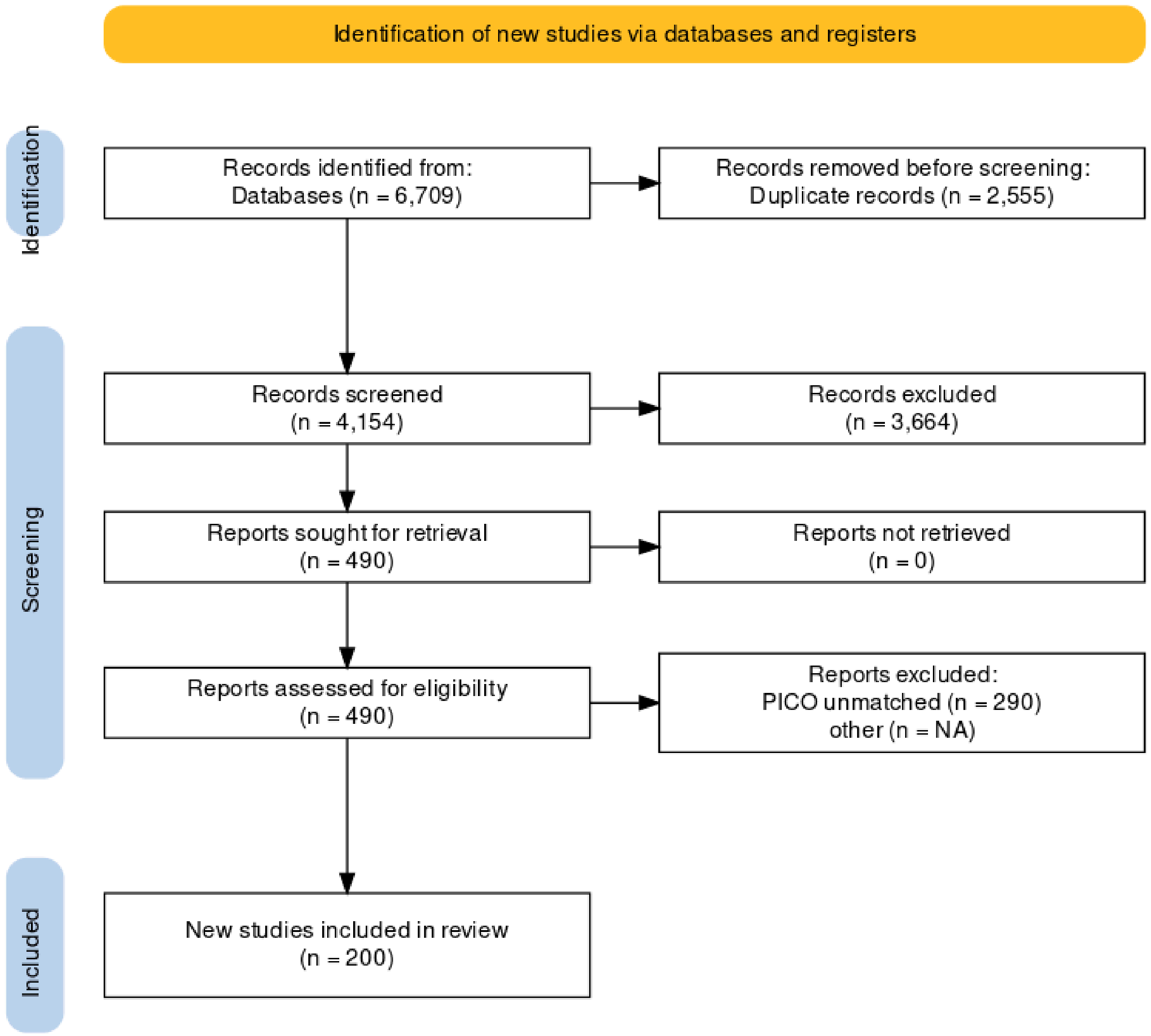


**Figure S1 -** PRISMA chart of the article selection process (data retrieved from Bonfanti et al. 2023). The diagram was produced with the online tool based on Haddaway et al. 2022.

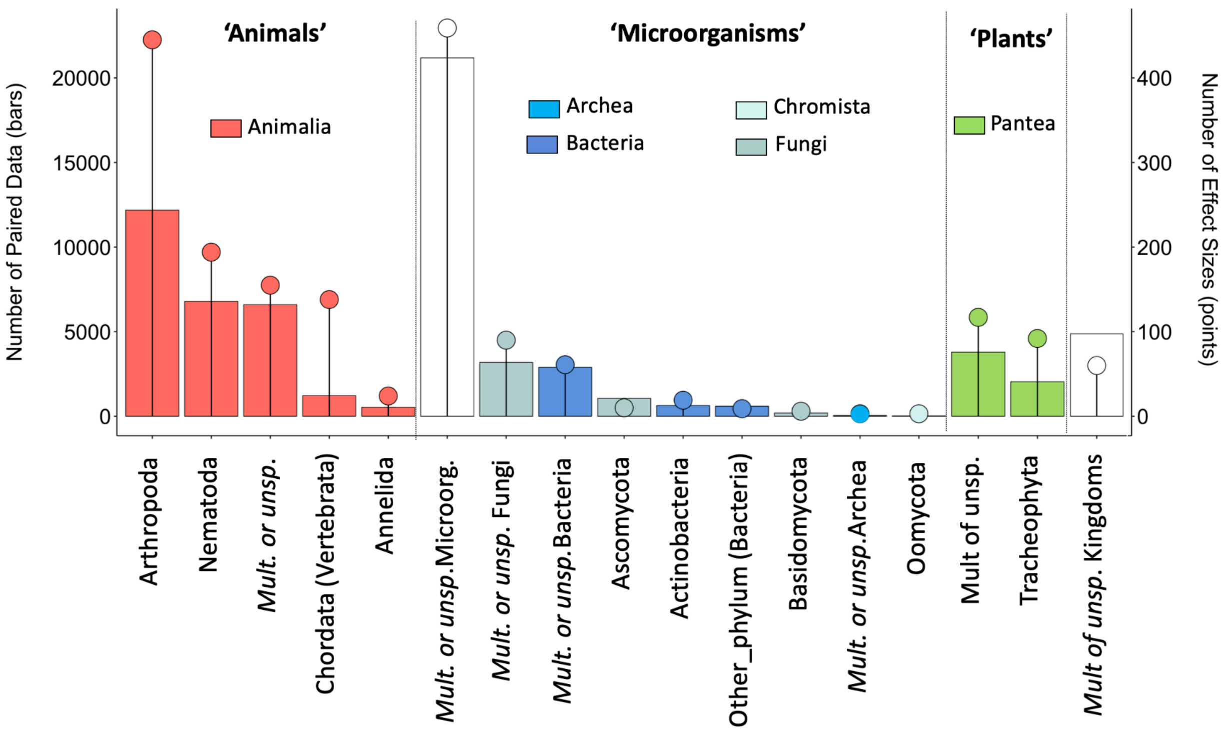


**Figure S2 -** The number of effect sizes (points: right axis), and paired data (bars: left axis) by broad taxonomic groups. Phylums are detailed in the x-axis. Mult.or.unsp.: multiple or unspecified, i.e. some effect sizes could not be attributed to a single taxonomic Phylum. Other_phylum (Bacteria): Acidobacteria, Bacteroidetes, Chloroflexi, Firmicutes, Proteobacteri. The colours represent the different Kingdoms. One meta-analysis may provide several effect sizes concerning different taxonomic groups.

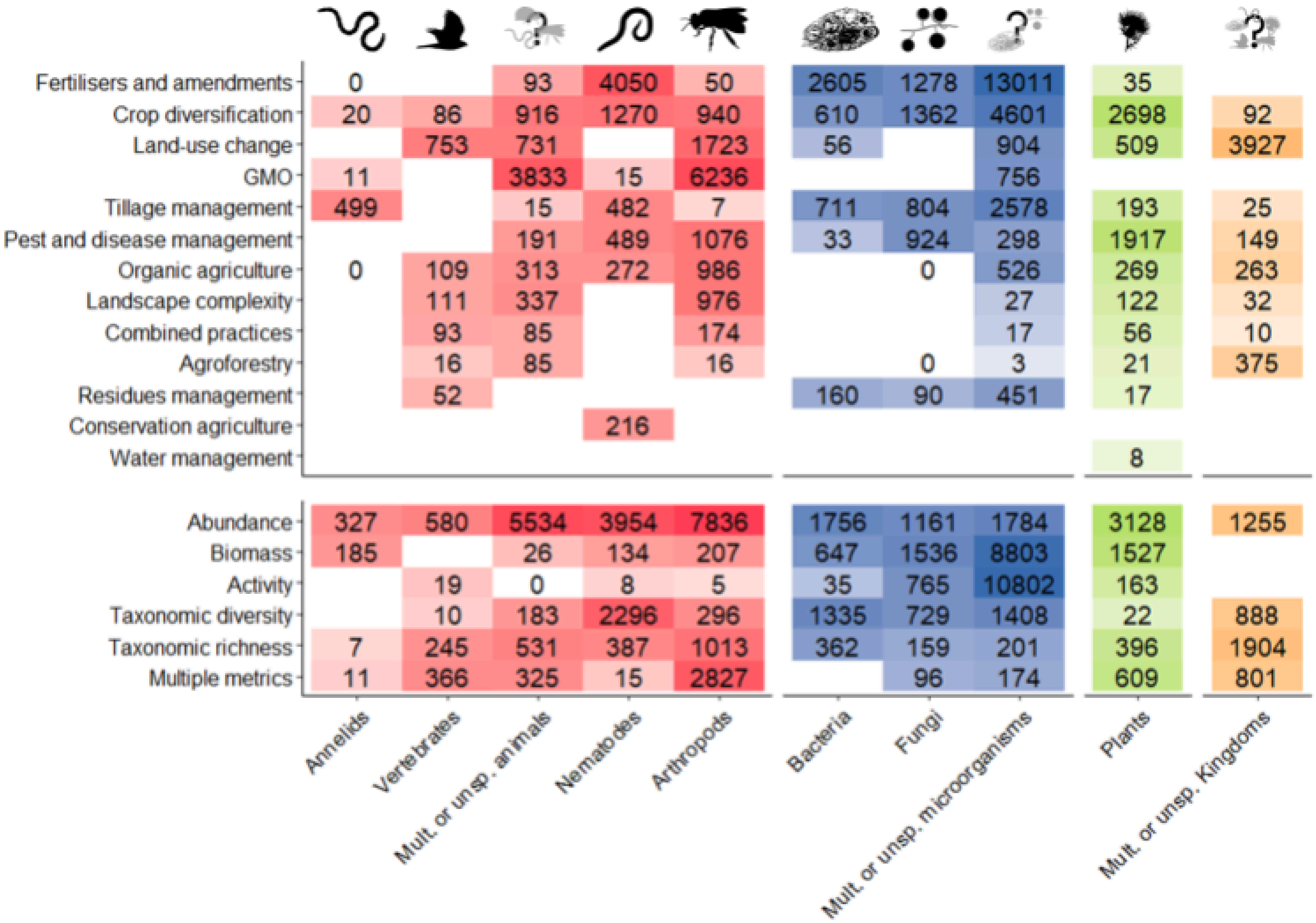


**Figure S3 -** Number of paired data per taxonomic group group, for each agricultural intervention and each biodiversity metric.

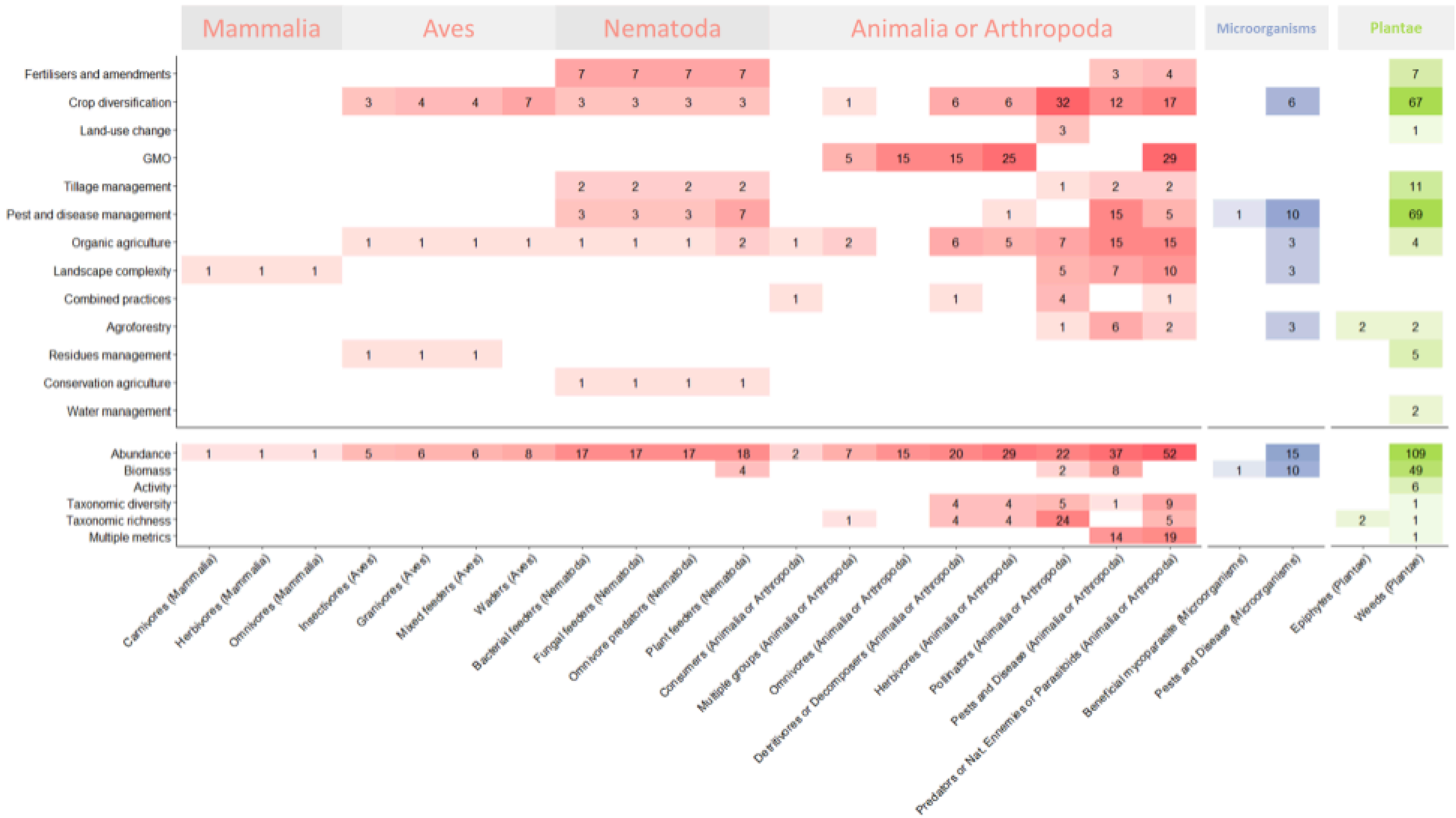


**Figure S4 -** Number of effect sizes per ecological group for each agricultural intervention and biodiversity metric

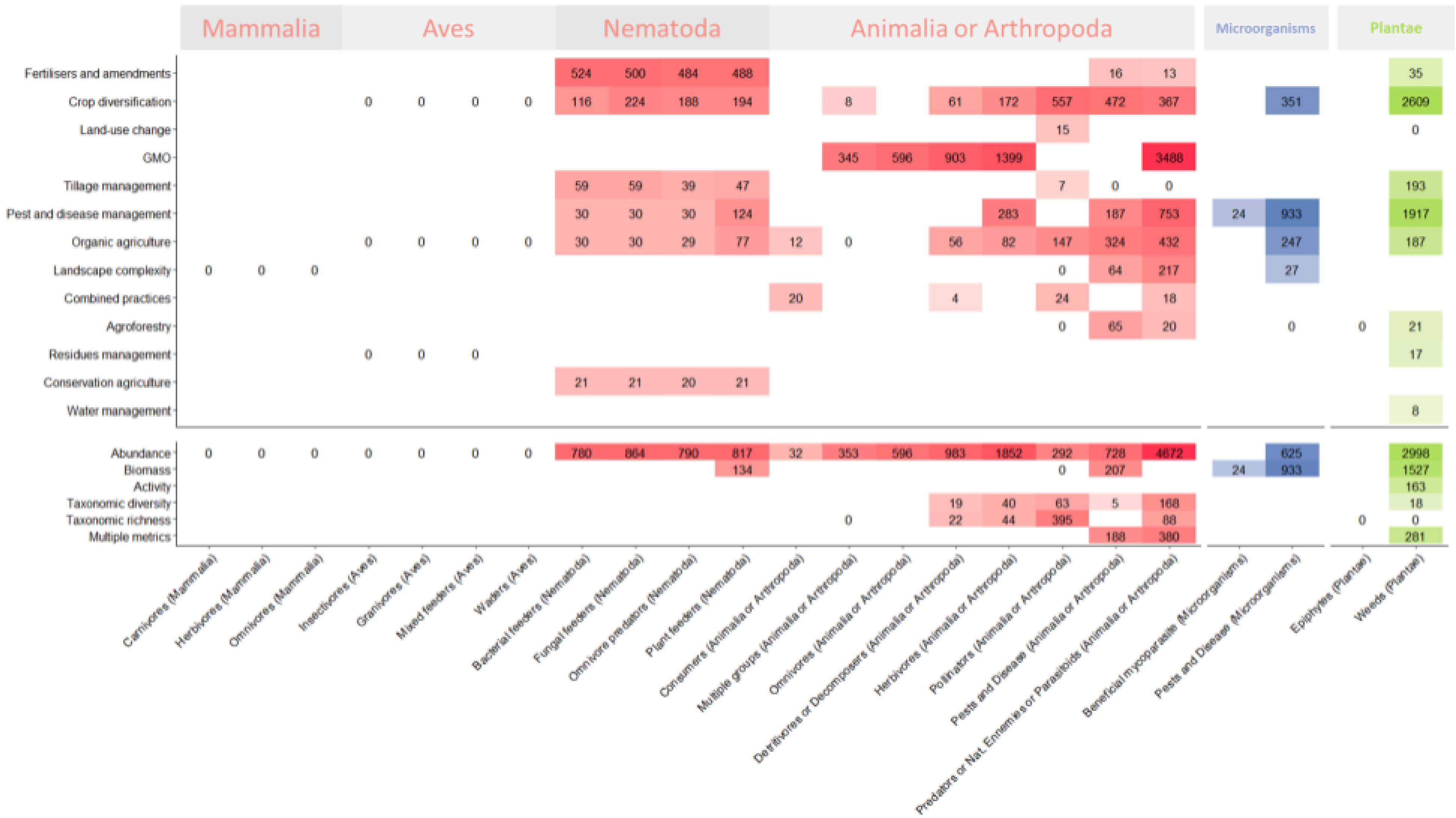


**Figure S5 -** Number of paired data per ecological group for each agricultural intervention and biodiversity metric

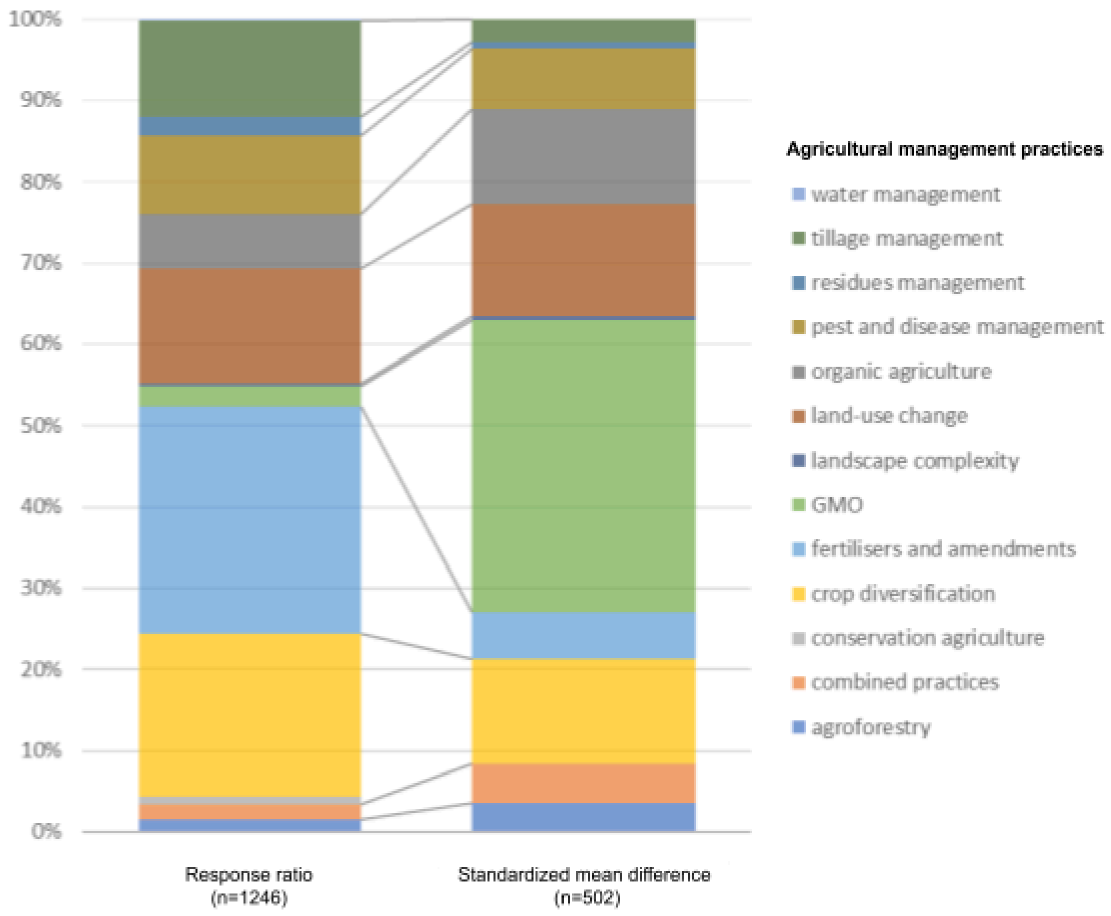


**Figure S6** - Stacked bar plots showing the two most represented effect size metrics categories (according to Borenstein 2009) in our database and their associated agricultural interventions.

N.B Note that bars are scaled to 100%, thus due to large differences in counts, bars' compartments of comparable size represent ca. 2.5x effect sizes for 'Response ratio' than for 'Standardised mean difference'.

**Table S3** - Raw data for Figure 6; percentage of occurrence of different taxonomic groups in our study and in Mammola et al 2023 study.

| ratio | Log Ratio | Taxonomic group | % occurrence in Mammola et al. | % occurrence in our study |
|---|---|---|---|---|
| 1,34788732 | 0,12965359 | Animalia | 0,37665782 | 0,507692308 |
| 1,06666667 | 0,02802872 | Annelida | 0,01193634 | 0,012732095 |
| 1,85416667 | 0,26814877 | Arthropoda | 0,12732095 | 0,236074271 |
| 0,42461538 | -0,37200427 | Chordata (Vertebrata) | 0,17241379 | 0,073209549 |
| 0 | -1 | Mollusca | 0,0066313 | 0 |
| 9,7 | 0,98677173 | Nematoda | 0,01061008 | 0,102917772 |
| 0 | -1 | Platyhelminthes | 0,00132626 | 0 |
| 0,4 | -0,39794001 | Archaea | 0,00397878 | 0,001591512 |
| 1,424 | 0,15350999 | Bacteria | 0,0331565 | 0,047214854 |
| NA | 1 | Chromista | 0 | 0,001591512 |
| 0,78518519 | -0,1050279 | Fungi | 0,07161804 | 0,056233422 |
| 0,36363636 | -0,43933269 | Mult. or unsp. Kingdoms | 0,08753316 | 0,031830239 |
| 0 | -1 | Algae | 0,0066313 | 0 |
| 0,26329114 | -0,57956376 | Plantae | 0,41909814 | 0,110344828 |
| 0 | -1 | Protozoa | 0,00132626 | 0 |
| 3 | 0,47712125 | Microorganisms | 0,11671088 | 0,350132626 |
